# Bacterial growth dynamics on a surface having a particulate antimicrobial agent

**DOI:** 10.1101/2024.05.30.596615

**Authors:** Cyrus Talebpour, Fereshteh Fani, Hossein Salimnia, Marc Ouellette, Houshang Alamdari

## Abstract

The morphological dynamics of microbial cell proliferation on an antimicrobial surface at an early growth stage was studied with *Escherichia coli* on the surface of a gel supplied with nanostructured AgNbO_3_ antimicrobial particles. We demonstrated an inhibitory surface concentration, analogous to minimum inhibitory concentration, beyond which the growth of colonies and formation of biofilm are inhibited. In contrast, at lower concentrations, colonies circumvent the antimicrobial activity of the particles and grow with a short lag time of a few hours. The applicability of these findings, in terms of estimating inhibitory surface concentration, was tested in the case of antimicrobial polymethyl methacrylate (PMMA) bone cement.

## 1. Introduction

The misuse of antibiotics has led to development of infections attributed to antimicrobial resistant (AMR) bacteria strains, which have been responsible for an annual 1.3 million deaths globally [1]. If nothing is done, this figure is projected to rise to 10 million by 2050 [2], with expenditures resulting in a global loss of $108 trillion USD [3]. The need to counter AMR may be achieved through strategies such as effectively reducing the microbial load on surfaces that come into contact with pathogenic microbial cells [4]. Any remedy for rendering antimicrobial properties to these surfaces is desired to present no systemic or localized toxicity, inhibit the proliferation of most species of microbial cells in contrast to a selected category of species, provide a robust system enabling ease-of-use, and involve a low production cost [5]. In this respect, silver coatings have been good candidates [6-8], thanks to their broad-spectrum antimicrobial activity through mechanisms such as production of reactive oxygen species (ROS) [9]. However, this approach is associated with drawbacks such as incompatibility of the conventional coating techniques with some important solids, altering the structure of surface, and corrosion with elution of silver ions, which ironically are the agent of antimicrobial activity [10].

Recently, we found a possible solution for the latter drawback by synthesizing nano-structured AgNbO_3_ nanoparticles, which, despite having diminished ion release rate, show strong antimicrobial activity [11]. This kind of antimicrobial activity, which we had defined as ‘through contact’, are associated with particulate agents and can be incorporated into the surface provided that the required amount of the particles is not below a threshold. Thus, a main question to be addressed for designing an antimicrobial surface employing these particles is introducing and quantifying a property akin to the minimum inhibitory concentration (MIC).

Another question, relevant for the antimicrobial surfaces in niche applications such as the interfacial surfaces of implants, is the formation and evolution of biofilms, i.e., structured communities of sessile bacteria [12] encased in a self-produced matrix of extracellular polymeric substances (EPS) [13]. In the context of AMR, the biofilm structure facilitates quorum sensing between cells responsible for altering gene expression, accumulating genetic material for cellular uptake, promoting horizontal gene transfer, and enabling less metabolically active cells known as persister cells to arise [14]. Up to 65-80% of all infections are associated with biofilm formation, much of which are implicated in chronic infections, in contrast to planktonic bacteria involved in acute processes [15]. Therefore, a phenomenological investigation of biofilm evolution on a surface having a concentration of antimicrobial particles sublethal to bacteria is a minimum necessity for actual implementation of an antimicrobial surface.

In this study we determine the surface minimum inhibitory concentration (SMIC) for surface-utilizing particles with antimicrobial activity through contact. The utility of these findings is illustrated in the case of polymethyl methacrylate (PMMA) bone cement. Furthermore, a qualitative description of colony formation and expansion at early stages are presented by time lapse microscopy on a model system, consisting of a gel having dispersed AgNbO_3_ nanoparticles on its surface. It is worth mentioning that incorporation of the right amount of AgNbO_3_ nanoparticles in PMMA bone cement is expected to confer permanent antibacterial property to the cement due to its extremely limited corrosion/dissolution rate, contrary to the incorporation of conventional antibiotics such as gentamicin.

## 2. Methodology

### 2.1 Preparation and characterization of nanostructured antimicrobial particles

Nanostructured AgNbO_3_ antimicrobial particles were synthesized using the Activated Reactive Synthesis (ARS) method as described previously [11, 16]. Briefly, the raw materials Ag_2_O (Sigma-Aldrich Corp) and Nb_2_O_5_ (Inframat^Ⓡ^ Advanced Materials LLC), with the weight ratio of 1 g to 1.147 g respectively (for each 1 g of Ag_2_O, 1.147 g of Nb_2_O_5_ powders), were mixed in a hardened steel crucible with high energy ball milling for 10 min. The mixture was transferred to a ceramic crucible and placed in an oven where it was gradually heated at a rate of 5 °C/min until the formation temperature, 1000 °C, was reached. The mixture was kept at this temperature for about 4 h and gradually cooled down at a rate of 10 °C/min to room temperature. The post-synthesis treatment included subjecting the powder to successive steps of high and low energy ball milling. The high energy ball milling was carried out using the 8000D Mixer/Mill^Ⓡ^ (SPEX SamplePrep, LLC). An 8001 hardened steel crucible and a milling media consisting of two balls of 12.70 mm and one ball of 6.35 mm in diameter were used for milling 7 g of material in each batch at an oscillation rate of 1060 cycles/min for a duration of 90 min. Low energy ball milling was carried out by transferring approximately 40 g of powder from the previous step (agglomerates) to a crucible containing hundreds of steel beads of 4.5 mm in diameter, which were made to rotate at 90 rpm by Szegvari Attritor System Type E Model 01-STD (Union Process, Inc.). To this, 10 mL of water was added and the attrition process was performed for 120 min. At the end of the operation the beads were rinsed with deionized water and the residue thus obtained was dried inside an oven with a temperature of 150 °C overnight to obtain the final nanostructured AgNbO_3_ particles. The particles were previously characterized in terms of size by the Malvern ZEN1600 Dynamic Light Scattering (DLS) machine and found to have an average size of 0.44 μm [11]. Density of the particles was measured using a AccuPyc II 1340 (Micromeritics Instruments Corp.) helium gas pycnometer. Powder samples of ∼ 8 g were weighted in a 10 mL stainless steel cell, and the density was obtained by dividing the mass of the sample to the volume measured by the pycnometer.

### 2.2 Bacterial strain and culture conditions

The bacterial strain used in this study was *Escherichia coli* ATCC # 25922 and was prepared as described below. Cell stock in 50% glycerol was taken from a -80 °C freezer and after thawing, 30-50 µL of it was transferred to a Tryptic Soy Agar (TSA) (with 5% sheep blood) plate (the P1 plate), inoculated by standard streaking, and incubated at 37 °C overnight under aerobic condition until colonies were visible. A well-formed representative colony from the plate was picked and inoculated into 3 mL of Tryptic Soy Broth (TSB) and incubated at 37 °C with 150 rpm shaking for 3 h. Then, a 1 mL culture was transferred to 2 mL autoclaved tubes and centrifuged at 8000 rpm for 8 min in a microcentrifuge. The harvested cells were resuspended in 1 mL TSB and 0.5 mL was used for OD_600_ measurement. Using an in-house OD_600_ vs cell count correlation database, a cell suspension of 10^8^ CFU/mL was prepared. The suspension was then serially diluted in TSB to a final concentration of 10^7^, 10^6^ and 10^5^ CFU/mL by pipetting and vortex mixing at each dilution step.

### 2.3 Planktonic growth curve and antimicrobial susceptibility

The growth of *Escherichia coli* ATCC # 25922 bacterial cells were determined using an automated incubation system (Biospa, BioTek) integrated with Cytation 5 multimode reader (BioTek). 10 μl of 1 × 10^7^ CFU/mL bacteria cells were inoculated in 990 μl of LB medium in the absence or presence of AgNbO_3_ nanoparticles at concentrations of 4, 8, 16 and 32 μg/ml in a Falcon^®^ 24-well plate. The plate was incubated at 37°C, and the OD_600_ was read at each 1 h for 20 h after shaking for 10 s using a Cytation 5 multimode reader. Data was processed using Gen5 software and reported with GraphPad Prism. The minimum inhibitory concentration (MIC) was determined as the concentration of particles for which no growth was observed after 24 h of incubation. All MICs were determined from at least three independent biological replicates.

### 2.4 Preparation of antimicrobial gel surfaces

5 mg of the nanostructured particles was weighted and resuspended in 5 mL of deionized water. The suspension was serially diluted to concentrations of 200, 100, and 50 μg/mL. To prepare an antimicrobial gel, standard A9 blood agar plate was left on the table for 30 min, then 5 μL of each particle suspension, after vortexing for 1 min, was dispensed at a spot in the first half of the plate. The procedure was repeated on the second half of the plate to obtain two technical replicates. On each half of the plate, 5 μL of water was dispensed on designated spots as control. The dispensed particle suspension spreads to a disk with a diameter of ∼ 8 mm and surface area of ∼ 50 mm^2^ before drying after 30 min. Thus, the particle density at the respective spots corresponding to 50 μg/mL, 100 μg/mL, and 200 μg/mL particle suspensions are respectively 5 ng/mm^2^, 10 ng/mm^2^, and 20 ng/mm^2^. Three biological replicates were prepared for each condition.

### 2.5 Seeding microbial cells on solid phase growth media and interrogating their growth

From a 1 × 10^5^ CFU/mL *Escherichia coli* ATCC # 25922 cell stock, 1 μL (nominally containing 100 CFU) was dispensed on each spot wherein particle suspensions and controls were previously dispensed. The dispensed cell stock spreads to a disk with a diameter of ∼ 4 mm and surface area of ∼ 12 mm^2^. The plates were placed inside the incubator and a rectangular part at the center of the spots, having an area of 2.05 mm^2^ (1.75 mm × 1.17 mm), were imaged with a metallurgical microscope (10× objective) in a bright and dark field at different time intervals. Each image represents approximately 1/6 of the entire spot area and the number of the growing colonies detected on it fairly well approximates 1/6 of the dispensed cells, verifying the homogenous dispersion of the bacteria cell on the surface. Using imageJ software, the microscopic images were analyzed for extracting the characteristic features of three representative colonies within one spot for each condition, and the average size of the 20 representative colonies from the two spots for each condition. Two other biological replicates for each condition were done for confirmation of the results obtained.

### 2.6 Preparing and testing antimicrobial disks

AgNbO_3_ loaded polymethylmethacrylate (PMMA) disks were prepared by mixing 10 g of the PMMA dry component (A formulation of 97% Poly(methyl methacrylate), Sigma-Aldrich, and 3% Benzoyl peroxide, Sigma-Aldrich) with different amounts of AgNbO_3_ (Control, 0.5%, 1%, and 2%) to obtain the desired W/W ratio. Then, 5 ml of liquid monomer (A formulation of 97% Methyl methacrylate, Sigma-Aldrich, and 3% N,N-Dimethyl-p-toluidine, thermo scientific) was added to the mixture and mixed until a homogenous paste was formed. This was immediately cast in a High-Density Polyethylene (HDPE) mold with an inner diameter of 12 mm and depth of 10 mm. After solidification, the disc was released from the mold and used in subsequent experiments. The naming conventions for the samples are hereafter designated as control PMMA and PMMA - X% AgNbO_3_ (X = 0.5, 1, 2).

The disks were tested for antibacterial properties following the CLSI M07 and CLSI M100 guidelines [17, 18]. Briefly, three replicates of each disk were sterilized by soaking in 70% isopropanol for one min, washing in sterile water for 2 min, and allowing them to air dry. The top surfaces of the disks were respectively loaded with a 50 μl of 1.5 × 10^6^ CFU/mL *Escherichia coli* ATCC # 25922 bacterial cell suspensions (corresponding to around 75000 bacterial cells) and, after allowing to dry in air for 3 h, were dropped in a tube of 5 mL TSB. 5 min after dropping the disks in TSB, a 1 mL sample from each growth media was transferred in a Falcon ® 24-well plate. The original tubes were incubated at 37 °C overnight. After incubation, the tube was checked visually for signs of growth (turbidity) or lack of growth (clear medium). For the plate, the OD_600_ was read each 1 h for 20 h after shaking for 10 s using a Cytation 5 multimode reader (BioTek).

## 3. Results and discussion

### 3.1 Planktonic growth dynamics

Fig 1 presents the planktonic growth dynamics, indicated by average optical density over three biological replicates versus incubation time. Based on the general shape of the curves, the impact of antimicrobial particles on the cell viability, being manifested in the form of prolonging the lag phase by 4 and 8 h, respectively for 4 and 8 μg/mL samples. This may be a manifestation of the “tolerance by lag” phenomenon, which appears to involve a generalized adaptive response to antibiotic stress [19]. Alternatively, one may attribute the prolonged lag phase to the presence of a subpopulation of antimicrobial persister cells in the entire inoculum 10^5^ CFU that have a phenotypic tolerance to the antimicrobial particles [20]. The second key observation is that the minimum inhibitory value is at most 16 μg/mL, as there is no sign of growth (increase in OD_600_) for this level of particle concentration after 20 h of incubation. Regarding these observations, some contrasts are expected with the case of exposing the microbial cells to antimicrobial particles on the surfaces for the following reasons. As the action of these particles is exerted via contact (since they do not release silver ions over the MIC level [11]) the growth inhibition on the surface has to occur either by incidental placing of a seeded cell at the proximity of a particle or via the cell motility leading to collisions.

**Fig 1.**
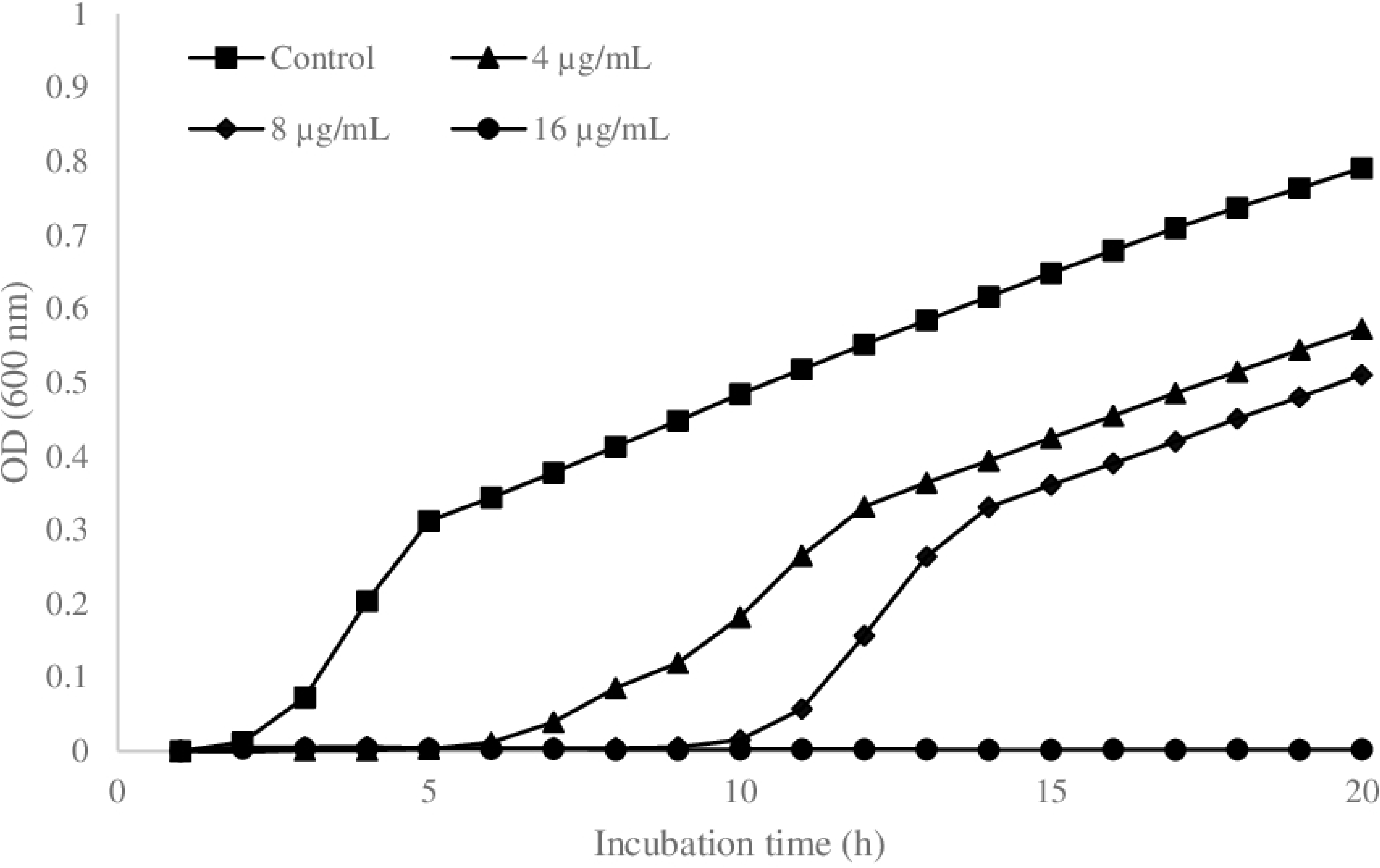
The growth curve of *Escherichia coli* ATCC # 25922 in LB broth in the presence of antimicrobial AgNbO_3_ particles with different concentrations. Each point is the average of three biological replicas.

### 3.2 Morphological and growth dynamics of colonies on a surface with no antimicrobial particles

The growth behavior of colonies on a plate with no antimicrobial particles performed as an independent test was selected as a baseline for assessing the impact of particles during the following experiments. The results are hereafter referred to as “trend”. The parameters selected for this purpose were colony shape and its overall size, which is defined as the diameter, *d*, of a disk with similar area, as specified in equation 1.

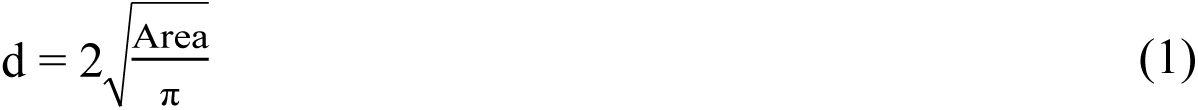

When 1 μL of a 1 × 10^2^ CFU/μL cell suspension was dispensed on the agar plate, 22 and 24 colonies were observed on the respective images taken from two spots. The area of the images of the two spots represented 1/6^th^ of the total seeded area. The average of these two numbers, i.e., 23, indicates the number of colonies in the examined area. The total number of the colonies can then be estimated as 6 × 23 = 138, which is close to the nominally expected number of the dispensed cells. Fig 2 presents zoomed images of a typical selected colony taken at different times of incubation of (2-5) h. The contours of the colonies at different time intervals are superimposed and presented at the center of the figure. The images may be used to describe the phases of colony morphology. In the first 3 h of incubation, the colony has expanded into an irregular shape, which, based on dark field images (not presented here), is in the form of a monolayer sheet of cells. At t = 3 h, the area of the colonies, averaged over the 10 colonies across the images of the two spots, is 391 μm^2^. After an extra hour of incubation, the average colony area increases to 2044 μm^2^, which is an increase by a factor of 5.2 (∼ 2.4 cycles). This factor is significantly smaller than the growth rate of 2.8 determined for microbial cells within a microcolony on an agar plate (See S1 Appendix). The deviation can be explained by noting that a second layer with a smaller area (presented by dashed green curve on the image corresponding to t = 4 h) was formed at the inner part of the first layer, thus increasing the total biomass of the colony by ∼ 0.4 cycles. Further incubation time increases these inner layers and the incremental change in colony’s apparent area drops to 3.6× (∼ 1.9 cycles). This indicates a transition from the early stage of single layer colony to the stage when the colony is in the form of an incomplete cone with many layers [21, 22]. Based on the published research in the literature, the described trend in colony morphology results from the bacterial motility [23-27] and formation of extracellular matrix and/or cellulose which might contain protein fibers [28].

**Fig 2.**
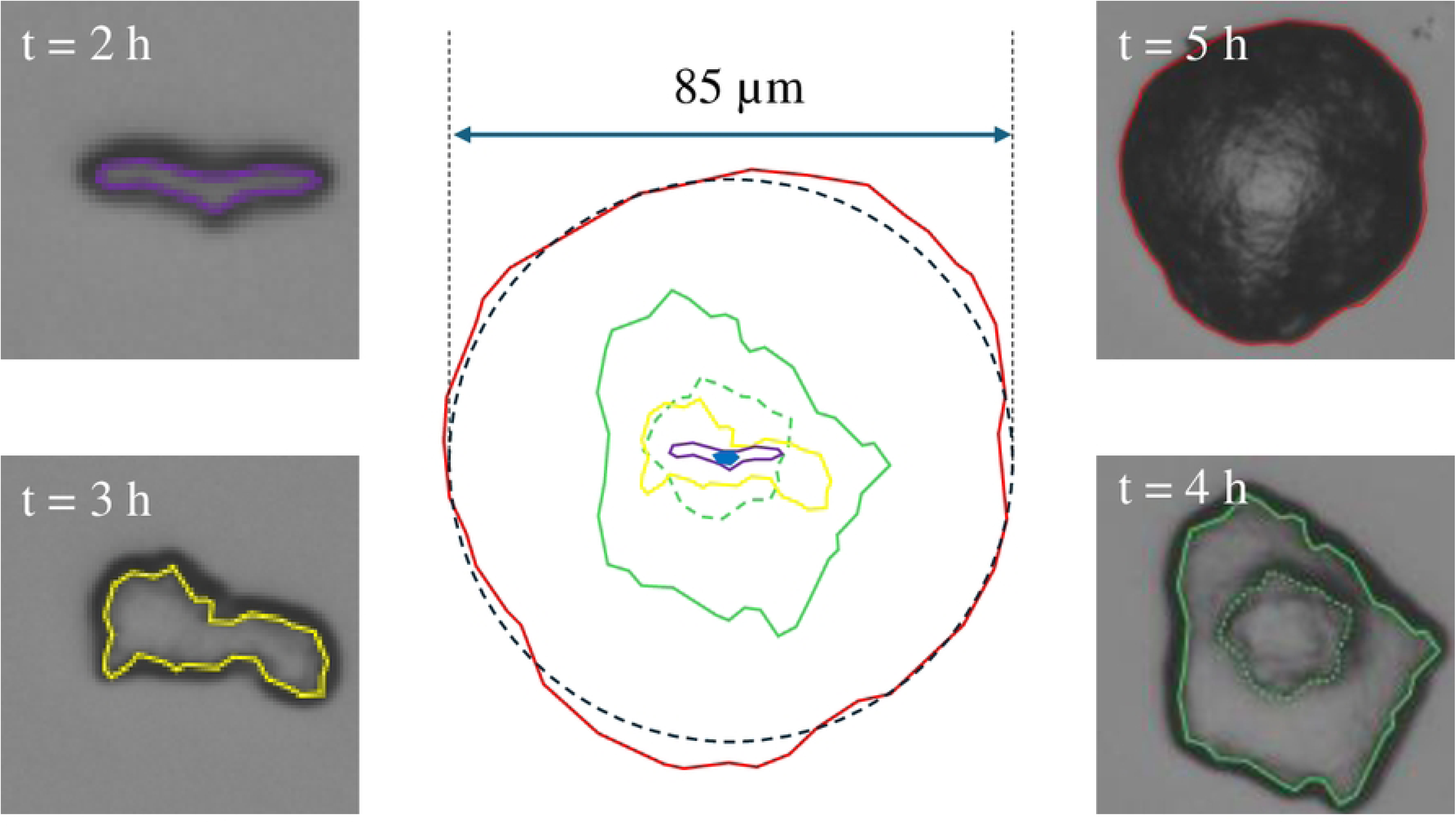
The morphological dynamics of three colonies grown on the surface of an agar gel and monitored with time lapse microscopy over 5 h of incubation. The contours indicate the colony boundary in the following order: Blue oval in center: Colony at t = 1 h, Violet: 2 h, Yellow: 3 h, Green: 4 h, and Red: 5 h.

This qualitative observation suggests a phenomenological description of a colony as schematically presented in Fig 3. The colony starts with a rugged monolayer cell as suggested by a single curved bright boundary in the accompanying dark field image. Then, distinct cell layers are added on the top of the first layer (z-direction). As the colony grows further, more layers are added and their boundaries in the dark-field image lose distinction. According to this model, the growth on solid phase takes place in 3 stages: 1) Early development phase, during which the colony is in the form of a single layer cell sheet and does not yet exhibit clear characteristic features, 2) Intermediate development phase, during which distinct cell layers are formed over the first layer, and, 3) Mature phase, when the layered structure makes a transition to a rough mass of cells. Based on this view, during the early development phase the cells are in direct contact with the surface and if the antimicrobial activity of the surface is not sufficient to prevent cell proliferation in a time scale of less than a few growth cycles, the upper layers may form. If the antimicrobial activity is exerted by contact, one can anticipate that these top layers be protected from the antimicrobial surface via the first layer, which acts as a barrier.

**Fig 3.**
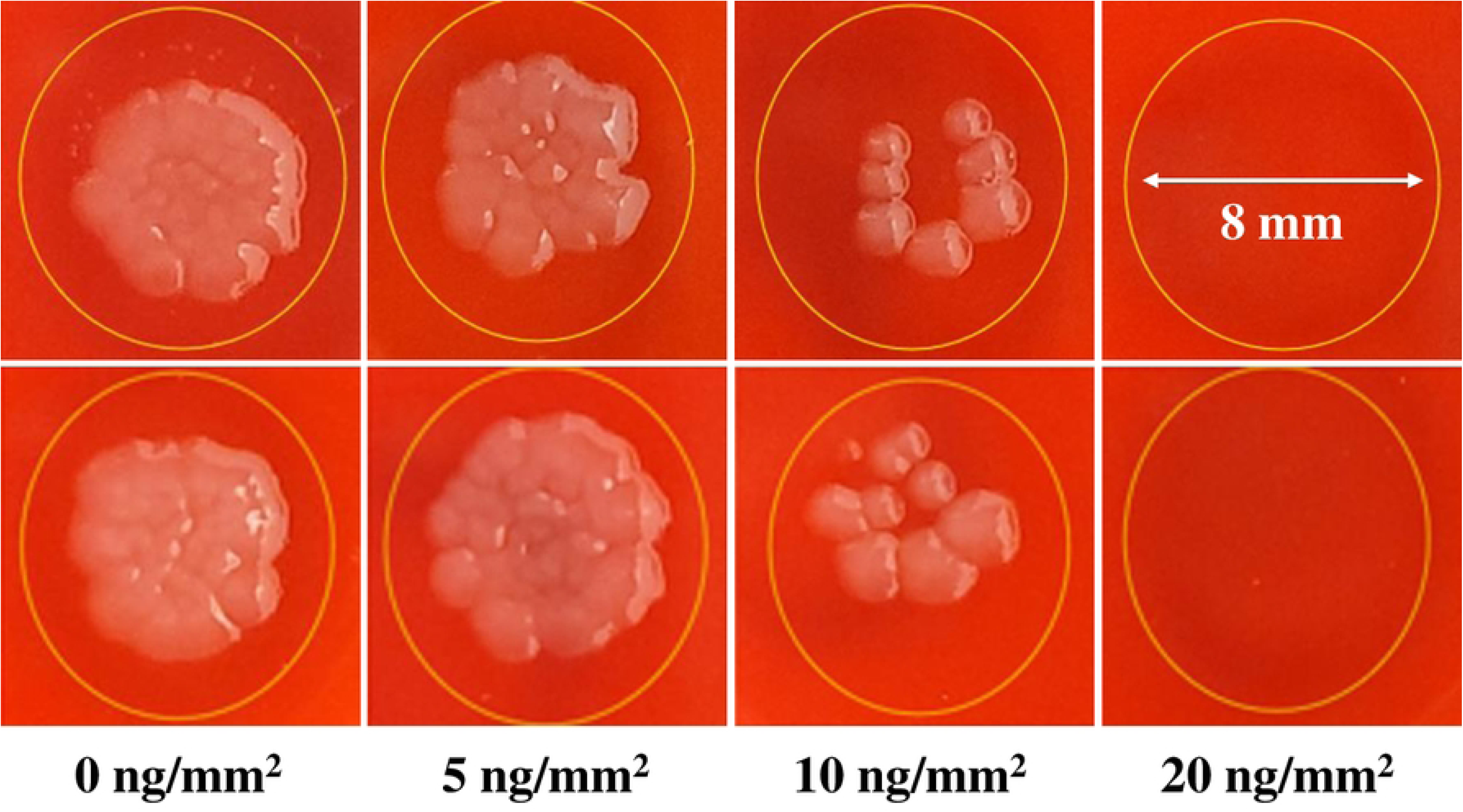
The schematic layered structure of a typical colony over incubation time. Schematics adapted from the reference [29], along with recorded dark-field images of a typical colony.

### 3.3 Growth dynamics of colonies on a surface containing antimicrobial particles

Fig A-D of S6 Appendix presents the partial image of the inoculated spot for different surface concentrations of antimicrobial particles of (0, 5, 10 and 20) ng/mm^2^ cases. The total number of colonies and the average size of 10 representative colonies grown on each of the two images for each condition after 6 h incubation were averaged and presented in Table 1. The images of the spots after leaving the plates at room temperature overnight, are presented in Fig 4.

**Fig 4.**
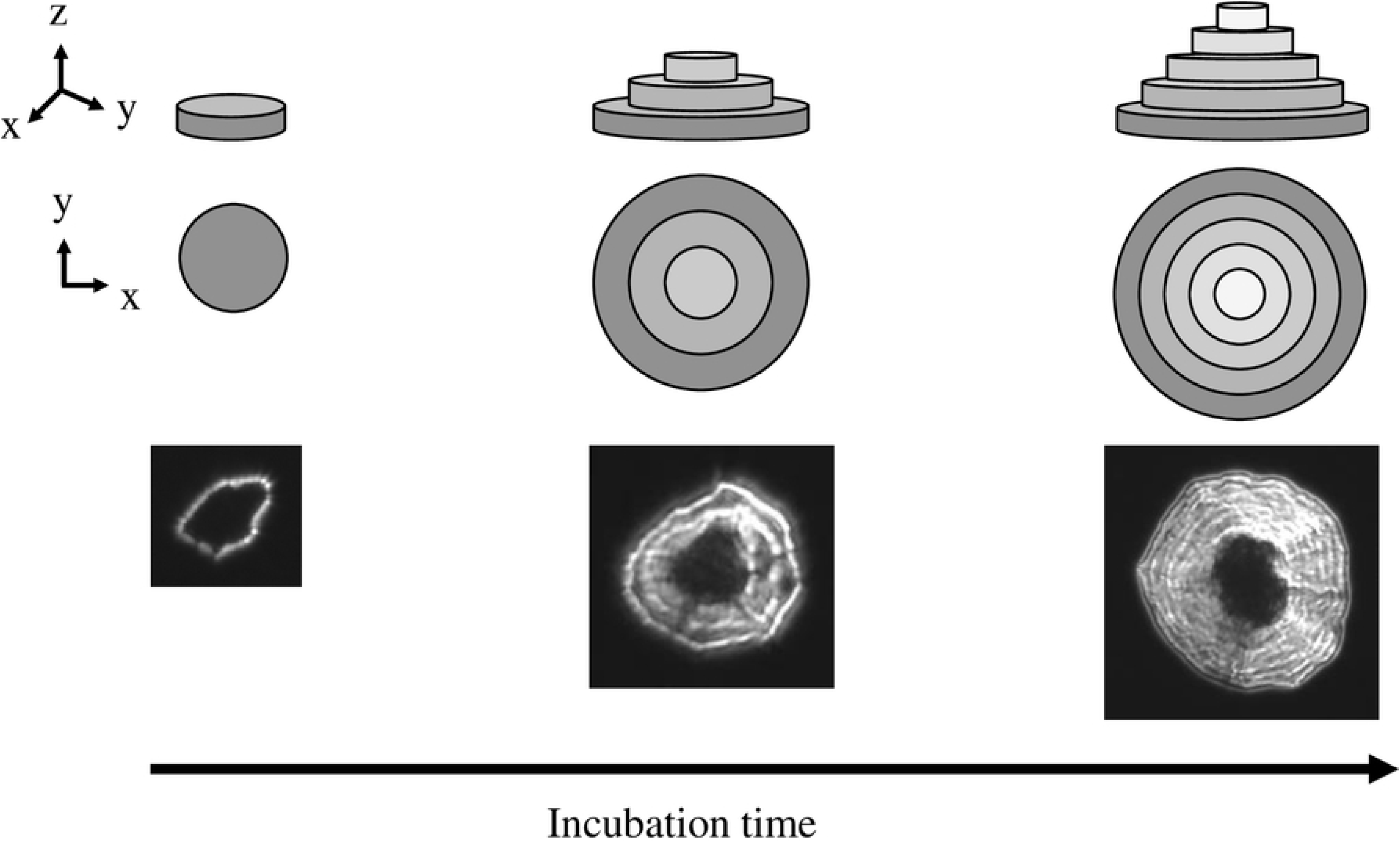
Shape of *Escherichia coli* colonies in two separate inocula on different BAP plates after leaving at room temperature overnight with varying concentrations of AgNbO_3_.

**Table 1.** The average number and sizes of colonies grown on 1.75 mm × 1.17 mm rectangular sections of two inoculated spots and their size distribution after 6 h incubation.

| Particle concentration (ng/mm<sup>2</sup>) | # of colonies | Average size (μm) |
| --- | --- | --- |
| 0 | 21 | 150 ± 16 |
| 5 | 18 | 77 ± 19 |
| 10 | 4 | 48 ± 14 |
| 20 | 0 | 0 |

The following observations were made: 1) 15% of the dispensed cells did not survive on the gel having 5 ng/mm^2^ of AgNbO_3_ particles. 2) The viability of the dispensed cells on the gel having 10 ng/mm^2^ of particles was 19%, and 3) There is no sign of growth for 20 ng/mm^2^. Accordingly, we took 20 ng/mm^2^ as equivalent to the MIC in the case of planktonic growth represented by the curves in Fig 1, albeit this does not imply an equivalency as it is the case for microdilution and agar dilution susceptibility tests for antibiotics [30]. The discrepancy results from the fact that the antimicrobial particles interact with bacterial cells not by diffusion but via contact.

Assuming an actual size of 0.69 μm for particles (See S3 Appendix), the surface MIC of 20 ng/mm^2^ translates to ∼ 2.7 mg/mL of AgNbO_3_ (See S4 Appendix); a value much higher than the 16 μg/mL obtained for planktonic cells (Fig 1). As it will be later seen, this observation is in good agreement with the exemplary case of loading a composite solid with the antimicrobial particles. However, first report our observation with the growth behavior of bacteria on the surface of gels containing sub-MIC surface concentration of antimicrobial particles.

In Fig 5, the size of microcolonies (average and standard deviation over 10 representative colonies) are presented as a function of time and were compared to those measured for the trend obtained in an independent test, as described in the previous subsection. Three observations can be made from this figure: 1) As it is expected, the growth behavior of the bacteria on a gel, as indicated by average colony size, is well repeated (control vs trend), 2) the lag time increases with concentration of antimicrobial particles. However, the lag time is smaller on surfaces than that in planktonic cases. For example, at ¼ and ½ the surface MIC, i.e. gels with 5 ng/mm^2^ and 10 ng/mm^2^ nanoparticles, the lag phases are 1 and 2 h respectively (Fig 1). In the planktonic cases, at antimicrobial concentrations of ¼ and ½ the MIC, the lag phases are 4 and 8 h respectively. 3) Finally, and similarly to planktonic cells (Fig 1), the surviving cells on the surface proliferate with an exponential growth rate, similar to the case of no antimicrobial particles (Fig 5).

**Fig 5.**
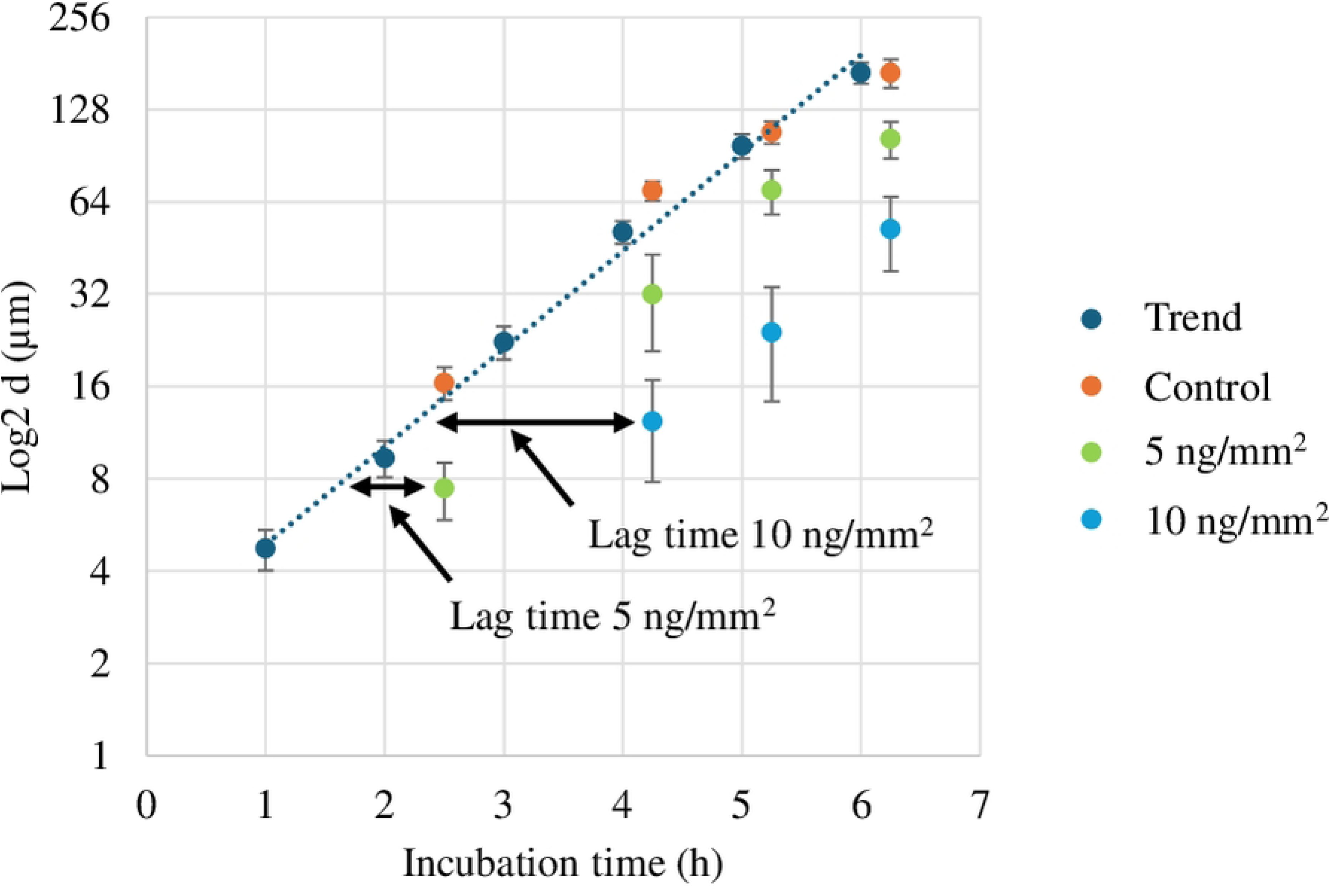
The average size of microbial cells grown on a gel surface having antimicrobial particles with surface densities of 5 and 10 ng/mm^2^ as compared to the size of colonies grown on a normal gel.

In Fig 6, the cropped microscopic images and time-lapse contours at 2 h, 4 h, 5 h, and 6 h of three colonies for control, 5 ng/mm^2^ and 10 ng/mm^2^ gels are presented. A dashed circle representing a radius of 150 μm, 100 μm and 70 μm was overlaid on the time-lapse contours of colonies from the control, 5 ng/mm^2^ and 10 ng/mm^2^ of AgNbO_3_ respectively. Within each of these dashed circles, the location of antimicrobial particles are presented as solid gray points. The original microscopic image of the gels after 5 h of incubation showing the location of the colonies are presented in Fig A-C in S6 Appendix.

**Fig 6.**
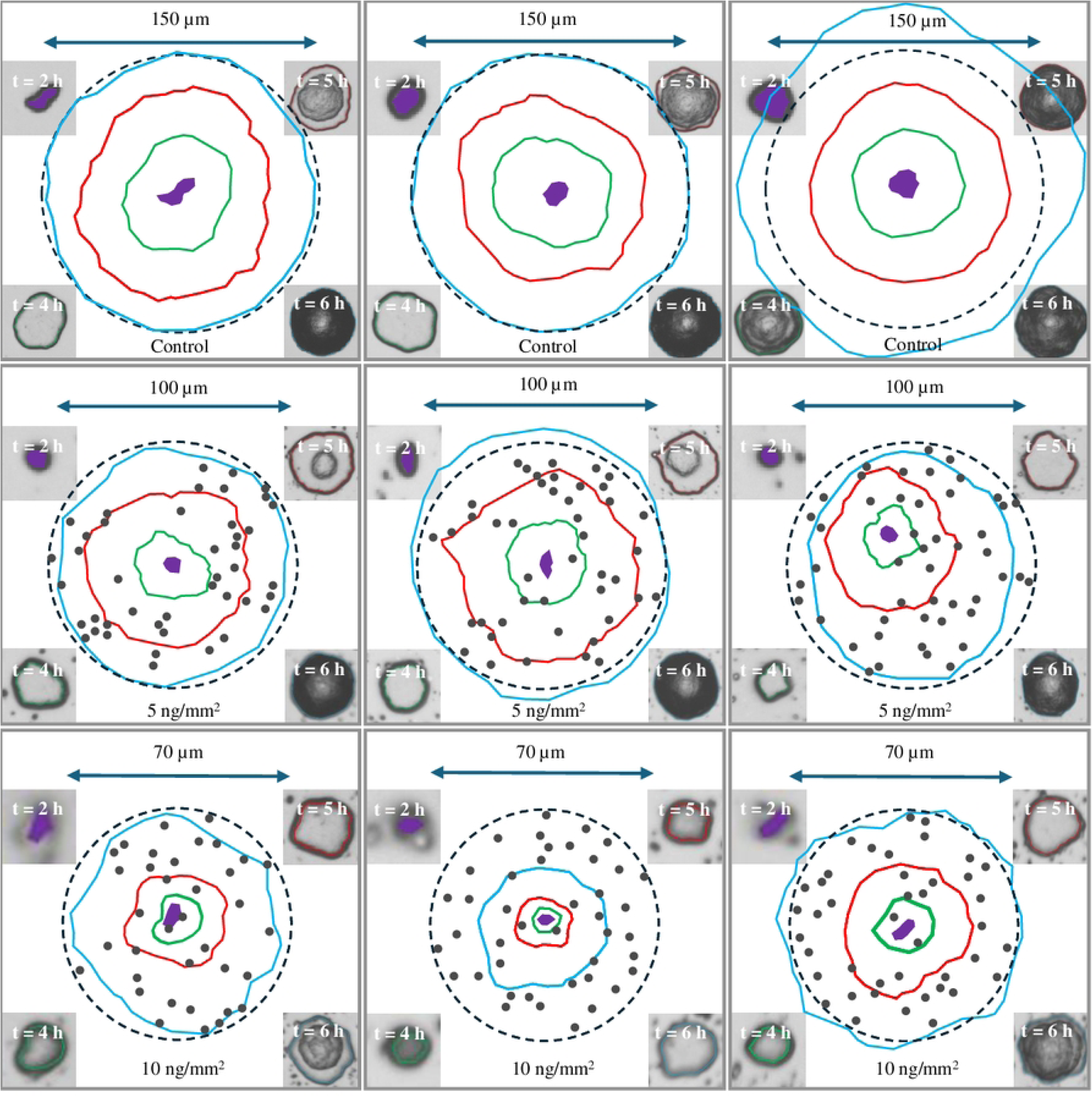
Evolution of the morphology of three representative colonies in the presence of different antimicrobial particle concentrations. First row: no antimicrobial particles, Second row: having antimicrobial particles with a surface density of 5 ng/mm^2^, and Third row: having antimicrobial particles with a surface density of 10 ng/mm^2^. Color code: Violet (2 h), Green (4 h), Red (5 h), and Cyan (6 h). The images of the colony after different incubation times are also shown at the corners in each case (starting from 2 h at upper left corner moving counterclockwise: 2 h, 4 h, 5 h, and 6 h). The antimicrobial particles are represented by solid gray points.

In the case of the control gel, similar to the phenomenological description of colonies in the independent baseline test, the colony starts with a cell monolayer and its size logarithmically increases. After 4 h of incubation, more cell layers are added over the first layer and finally the colony matures by the formation of a rough biomass structure. For the case of growth on gels with antimicrobial particles, the general trend is similar to the control, except the events have been delayed by lag time. This implies that once the seeded cell is able to overcome the antimicrobial induced lag phase, it is able to proceed with proliferation as part of a biofilm.

We hypothesize that just after being seeded, the cell starts preparing for the exponential growth phase by trying to accumulate transient metals [31]. The presence of a silver-containing compound in the vicinity perturbs this process. However, if a cell could overcome this hurdle and successfully divide, its descendants produce EPS, which acts as a structural barrier and provides tolerance to the antimicrobial particles [32]. In other words, the surface density of the particles must be sufficient enough to prevent the proliferation from the beginning, otherwise bacteria will create a protective barrier and continue to grow. This finding is of special importance in designing appropriate antibacterial surfaces, namely the bone cements which is our main goal. The bone cement must have sufficient particle density at its surface to prevent the proliferation of bacteria cells but not too high to be safe for human cells.

### 3.4 Bacterial proliferation on the surface of a composite solid loaded with antimicrobial particles

The predictive capability of our findings on the gel surface MIC was tested by assessing the antimicrobial level of a composite material. In this regard, we used PMMA as a solid, which is widely used as bone cement in orthopedic operations. The PMMA disks, loaded by different amounts of AgNbO_3_ particles, were inoculated with ∼ 75000 CFU of *Escherichia coli* and the survival of the cells were assessed. The result is summarized in Fig 7. The first observation is the 3 h of lag in the case of the PMMA - 0.5% AgNbO_3_ bone cement. This is more similar to the ∼ 2 h lag for the 10 ng/mm^2^ gel (as indicated in Fig 5) than to the ∼ 8 h of lag near MIC during the planktonic growth case of Fig 1. The second key observation is related to the surface MIC at the PMMA - 1% AgNbO_3_. According to the data presented in S2 Appendix, the estimated surface density of the AgNbO_3_ particles for 20 ng/mm^2^ gel is 1.93 × 10^4^ particles/mm^2^. If the particles are loaded in the bulk gel, this would correspond to an equivalent volume of 2.68 × 10^-3^ g/mL loading (See S4 Appendix). The mass density of PMMA is 1.18 g/mL [33]. If the bone cement surface behaved exactly similar to the particle loaded gels, 2.68 × 10^-3^ g/mL loading would correspond to (2.68 × 10^-3^ g/1.18g) × 100% = 0.23% of the cement, in w/w. The value is indeed not far from the observed 1% w/w for the particle loaded PMMA when the impact of inhomogeneous particle distribution on the surface is taken into consideration. More details on this respect are provided in S7 Appendix.

**Fig 7.**
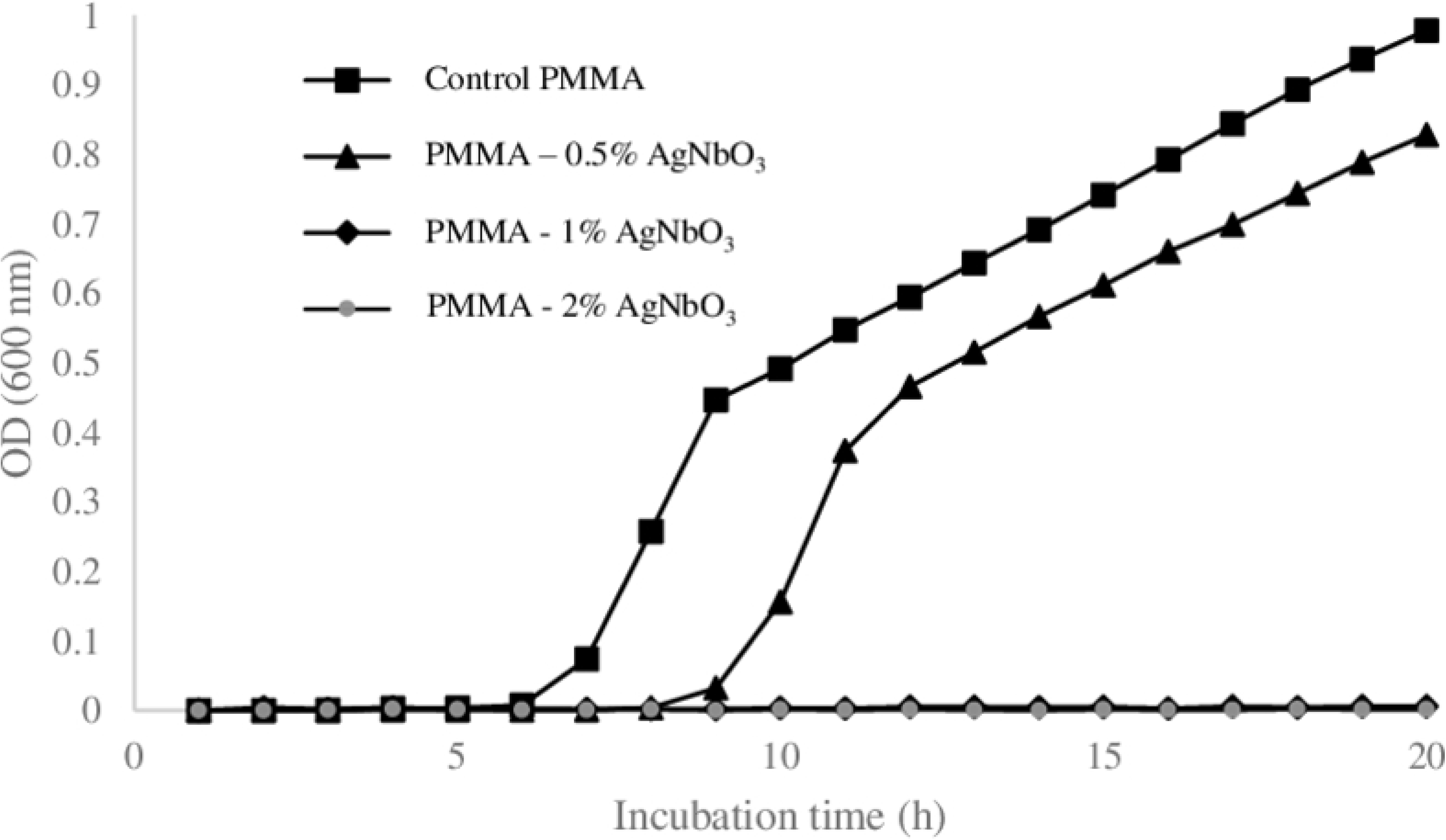
The optical density of the broth in which the *Escherichia coli* ATCC # 25922 cells contacted with the surfaces of PMMA samples, having different amounts of AgNbO_3_, are incubated. Each data point is average over 3 biological replicas.

Another factor, which contributes to the minor discordance between the antimicrobial efficacy of the antimicrobial particles on gel surface and those loaded into the bone cement, is the dissimilarity of inoculum in the two cases. We used fewer cells on the gel to avoid crowding which causes the merging of the colonies. On the other hand, we used a higher amount of cells on the cement surface to increase our confidence in calling the surface a true antimicrobial. Nonetheless we believe that the experiments on the gel are a good proxy for the microbial proliferation on the antimicrobial surfaces produced by loading antimicrobial particles.

## Conclusion

In conclusion, the growth of microbial colonies on the surface of a gel loaded with antimicrobial nanostructured AgNbO_3_ particles was investigated. It was found that similar to the planktonic case, the impact of sublethal levels of the antimicrobial particles is manifested as prolonged lag phase and a corresponding surface minimum inhibitory concentration (SMIC) could be defined above which the bacterial cells do not form colonies. The SMIC obtained in this way is a good estimator for the required antimicrobial concentration to be loaded into a solid to provide its surface with the antimicrobial property.

## Supporting information

S1 Appendix. Measuring solid phase growth rate

S2 Appendix. Surface density of AgNbO_3_ particles on antimicrobial gels S3 Appendix. Estimating average particle size, volume and mass

S4 Appendix. Determining the equivalent MIC

S5 Appendix. Determining distance between particles S6 Appendix. Supplementary information for Fig 6

S7 Appendix. Size distribution and density of the antimicrobial particles on the bone cement

